# Evolution under low antibiotic concentrations: a risk for the selection of *Pseudomonas aeruginosa* multidrug resistant mutants in nature

**DOI:** 10.1101/2021.04.21.440750

**Authors:** Fernando Sanz-García, Sara Hernando-Amado, José Luis Martínez

## Abstract

**BACKGROUND:** Antibiotic pollution of non-clinical environments might have a relevant impact on human health if resistant pathogens are selected. However, this potential risk is often overlooked, since drug concentrations in nature are usually below their minimal inhibitory concentrations (MICs). Albeit, antibiotic resistant bacteria can be selected even at sub-MIC concentrations, in a range that is dubbed the sub-MIC selective window, which depends on both the antibiotic and the pathogen.

**OBJECTIVES:** Determine the sub-MIC selective windows of seven antibiotics of clinical relevance in the opportunistic pathogen *Pseudomonas aeruginosa* and evaluate the risk for selecting resistant mutants in nature, based on published data about the amount of antimicrobials detected in natural environments.

**METHODS:** We conducted evolution experiments of *P. aeruginosa* PA14 in presence of sub-MIC concentrations of ceftazidime, amikacin, levofloxacin, ciprofloxacin, tetracycline, polymyxin B or imipenem, and measured drug susceptibility of the evolved populations.

**RESULTS:** Sub-MIC selective window of quinolones was the largest, and the ones of polymyxin B and imipenem, the narrowest. Clinically relevant multidrug resistant (MDR) mutants (presenting MICs above EUCAST clinical breakpoints) arose within the sub-MIC selective windows of the majority of antibiotics tested, being these phenotypes probably mediated by efflux pumps′ activity.

**DISCUSSION:** Our data show that the concentration of antibiotics reported in aquatic ecosystems -colonizable by *P. aeruginosa*- are, in occasions, higher than the ones able to select MDR mutants. This finding has implications for understanding the role of different ecosystems and conditions in the emergence of antibiotic resistance from a One-Health point of view. Further, it highlights the importance of delineating the sub-MIC selective windows for drugs of clinical value in pathogens with environmental niches, in order to evaluate the health risks due to antibiotic pollution of natural ecosystems and ultimately tackle antibiotic resistance.

## INTRODUCTION

Selection of antibiotic resistant mutants by an antibiotic takes place just in a range of concentrations. It had been generally accepted that this selective window (Zhao and Drlica 2002) spans from the minimal inhibitory concentration (MIC) for a wild-type strain to the mutant preventive concentration (MPC), which hinders the growth of single-step resistant mutants (Gullberg et al. 2011). However, it has been recently evidenced that selection of *de novo* resistant mutants, even highly resistant ones, may also occur under sub-MIC antibiotic concentrations (Wistrand-Yuen et al. 2018), which may pollute all kind of clinical and non-clinical environments (Danner et al. 2019). Besides their role in selection of resistant mutants, these concentrations present a non-negligible effect on the evolution of antibiotic resistance (AR) of bacterial populations, since they may modify mutation rates, biofilm formation and horizontal gene transfer (HGT), among other traits that relate to emergence and dissemination of AR (Gutierrez et al. 2013; Laureti et al. 2013; Romero et al. 2011). It is worth mentioning that the lowest antibiotic concentration able to select resistance, termed the minimal selective concentration (MSC), may drastically differ between antibiotics and bacteria (Gullberg et al. 2011; Gullberg et al. 2014; Sanz-Garcia et al. 2020). Therefore, the extent to which MSCs of distinct drugs could boost AR development in human pathogens, merit to be extensively delved into; a subject that has not been sufficiently researched yet (Gullberg et al. 2014; Lundstrom et al. 2016; Murray et al. 2018; Oz et al. 2014; Stanton et al. 2020).

This topic takes primary importance in the case of microorganisms with an environmental origin, because they can encounter sub-MIC antibiotic concentrations not only in clinics (i. e., due to limited drug accessibility, low-dose prophylactic therapies, poor patient adherence, as an outcome of incomplete dosing, or by the use of substandard medicines that often lack the stated amount of active pharmaceutical ingredient -API-) (Ching and Zaman 2020; Fisher et al. 2018; Kelesidis and Falagas 2015); but also in a great number of natural environments (lakes, rivers, wastewater treatment effluents, sludge, soils or even drinking water), mainly ascribed to agricultural activities and wastewater treatment (Danner et al. 2019; Martinez 2008; Wei et al. 2012). In fact, the data on the amount of different antimicrobials in diverse environments around the globe is rather concerning. In particular, 60.79 mg/l of tetracycline in a wastewater treatment plant in Northern China (Hou et al. 2016), 31 mg/l of ciprofloxacin in a pharmaceutical effluent in India (Larsson et al. 2007), 484 µg/l of oxytetracycline in a Chinese river (Li et al. 2008), or 53.8 µg/l of sulfamethoxazole in Mozambique surface waters (Segura et al. 2015), have been reported. From a One-Health angle, the interconnection between these environmental habitats and contiguous ecosystems could entail a critical risk to human health, since resistant bacteria selected under low drug concentrations may conceivably spread to human hosts (Hernando-Amado et al. 2019).

In this respect, one of the microorganisms that arises utmost concern is *Pseudomonas aeruginosa*, a free-living bacterium that is ubiquitous in nature -including aquatic domains-, thanks to its metabolic versatility (Crone et al. 2020; Mena and Gerba 2009). This opportunistic pathogen is one of the main causes of lung and airway infections in hospitalized patients; additionally, it is the major artificer of morbidity and mortality in patients suffering from cystic fibrosis and chronic obstructive pulmonary disease (Martinez-Solano et al. 2008; Talwalkar and Murray 2016). One of the reasons behind the threat to human health this bacterial species represents is its low susceptibility to several drugs (Alvarez-Ortega et al. 2011), as well as its ability to achieve resistance by acquiring chromosomal mutations (Lopez-Causape et al. 2018), among other mechanisms. In particular, multidrug (MDR) efflux pumps are likely one of the most outstanding sources of AR in *P. aeruginosa*, either providing intrinsic resistance or conferring mutational acquired one, via mutations that result in the overexpression of these systems and a heightened extrusion of their substrates (Camilo Barbosa et al. 2021; Poole 2001). MexAB-OprM, MexCD-OprJ, MexEF-OprN and MexXY-OprM are efflux pumps of the Resistance Nodulation Division (RND) family with a leading role in *P. aeruginosa* infections (Alcalde-Rico et al. 2016), since they may concurrently turn futile the action of various drugs that are usually applied to manage these infections, as β-lactams, aminoglycosides or quinolones (Bassetti et al. 2018; Masuda et al. 2000b). Thus, predicting the range of sub-MIC concentrations of clinically relevant drugs that might select for AR in *P. aeruginosa*, and the level of resistance and cross-resistance the selected mutants could acquire, is compelling; especially for uncovering the real risk associated with the presence of mild antibiotic concentrations in both clinical and natural ecosystems.

Herein, by using adaptive laboratory evolution (ALE) assays (Bouma and Lenski 1988) with a range of sub-MIC drug concentrations, we define the MSC and, consequently, the width of the sub-MIC selective window of 7 antibiotics of interest for *P. aeruginosa* therapies. Furthermore, we analyze if *de novo* selected antibiotic resistant mutants display cross-resistance or collateral sensitivity to antimicrobial agents used in clinical settings, and examine the role MDR efflux pumps might play in these phenotypes. Finally, based on our results and former reports, we point to the current risk of selecting AR in clinical and non-clinical circumstances.

## METHODS

### Bacterial growth measurement

Growth curves of *P. aeruginosa* PA14 in the presence of 1/100, 1/50, 1/25, 1/10, 1/5 and 1/2 (plus 1/200, in the case of ciprofloxacin) of its MIC to 7 antibiotics (ceftazidime, amikacin, levofloxacin, ciprofloxacin, tetracycline, polymyxin B and imipenem) were ascertained. A 10 µl sample of an overnight culture of this strain was inoculated in 140 µl of Luria Bertani (LB, Pronadisa) or Mueller Hinton II (MH II, Pronadisa) medium (in the case of polymyxin B) in presence of each concentration and in absence of antimicrobial, to a final OD_600_ of 0,01, in 96-well microtitre plates (Nunclon_TM_ Delta Surface; ThermoFisher Scientific, Waltham, MA, USA). Growth (OD_600_) from three independent replicates was monitored every 10 minutes by an Infinite 200 Plate Reader (TECAN, Männedorf, Switzerland) for 30 hours at 37 °C, with 20 seconds of shaking before each measurement.

### Short-term ALE assays

One hundred and forty four bacterial populations from a stock culture of *P. aeruginosa* PA14 were subjected to a 9 days ALE in presence of sub-MIC concentrations of ceftazidime, amikacin, levofloxacin, ciprofloxacin, tetracycline, polymyxin B or imipenem, ranging from 1/200 to 1/2 of MIC of the parental strain; being ceftazidime MIC = 4 µg/ml, amikacin MIC = 4 µg/ml, levofloxacin MIC = 4 µg/ml, ciprofloxacin MIC = 0,5 µg/ml, tetracycline MIC = 15 µg/ml, polymyxin B MIC = 4 µg/ml and imipenem MIC = 0,5 µg/ml. Cultures were grown in LB or MH II (in the case of polymyxin B) at 37 °C, with shaking at 250 rpm. Four replicates were submitted in presence of each drug concentration and without antibiotic. Each day, the cultures were diluted (1/100) in fresh medium, adding 10 µl of bacteria in 1 ml of LB or MH II, either containing the corresponding sub-MIC drug concentration or in absence of antibiotic. All replicate populations were preserved at -80 °C at 3, 6 and 9 days of each ALE assay.

### Antibiotic susceptibility assays

In order to monitor susceptibility changes along the ALE assays, the MICs to the selective antibiotics of the evolved populations were ascertained every 3 days, by using MIC test strips (Liofilchem®) at 37 °C in MH or MH II (in the case of polymyxin B) agar plates. The susceptibility to a broad range of antibiotics: ceftazidime, amikacin, levofloxacin, ciprofloxacin, tetracycline, polymyxin B, imipenem, aztreonam, fosfomycin, chloramphenicol and erythromycin was determined in the final evolved populations in which an increased resistance level (≥2-fold of the parental strain MIC value) to the selective antibiotic had been observed. When required, MICs in the presence of 25 µg/ml of Phe-Arg β-naphthylamide dihydrochloride (PAβN) (Merck) were also determined.

### Determination of β-lactamase activity

The *P. aeruginosa* populations selected after 9 days of evolution, as well as their parental strain PA14, were grown overnight at 37 °C and 250 rpm in 20 ml of LB medium. Next, they were spun down by centrifugation (10 min at 7000 rpm) and resuspended in 500 µl of 0,1 M Na_2_HPO_4_ (pH 7,4) buffer. Sonication (0,7 Hz) on ice and centrifugation (15 min at 13000 rpm and 4°C) were performed on these solutions to prepare crude protein extracts. Their protein content was quantified by using Bradford protein assay, with bovine serum albumin as standard. Afterwards, β-lactamase activity was measured as the change in absorbance (486 nm) of the chromogenic cephalosporin nitrocefin at 500 µg/ml (Oxoid, Basingstoke, United Kingdom), which was added to each sample. 0,1 M Na_2_HPO_4_ (pH 7,4) was used as the test buffer. This absorbance quantification was carried out using a Spark 10M Plate Reader (TECAN, Männedorf, Switzerland) for 2 hours at 37 °C, with measurements every 2 minutes.

## RESULTS

In the current work, we delimited the *P. aeruginosa*’s sub-MIC selective windows for 7 antibiotics. Ceftazidime, amikacin, levofloxacin, ciprofloxacin, polymyxin B and imipenem were chosen as representative members of the separate structural families that are regularly used for treating infections caused by this pathogen (Bassetti et al. 2018). Two quinolones were included in light of their foreseeably low MSC (Gullberg et al. 2011), which would be of significant interest. Although *P. aeruginosa* is intrinsically resistant to tetracycline (Morita et al. 2001), thence this antimicrobial is not of choice when it comes to therapies, prior works have revealed this antibiotic to be present in substantial amounts in natural habitats (Hou et al. 2016; Ok et al. 2011; Rodriguez-Mozaz et al. 2020). In addition, there exists evidence of substandard tetracycline usage - which does not have the stated quantity of API-in developing countries (Johnston and Holt 2014; Tabernero et al. 2019). We have previously shown that tigecycline, an antibiotic to which *P. aeruginosa* is intrinsically resistant too, selects for mutants displaying cross-resistance to clinically useful antibiotics (Sanz-Garcia et al. 2018b). Consequently, it is relevant to examine if sub-MIC tetracycline concentrations that could be found in natural ecosystems might select cross-resistant mutants to other antipseudomonal drugs, even though this antibiotic is not used for treating *P. aeruginosa* infections.

### Effect of sub-MIC antibiotic concentrations in *P. aeruginosa* fitness

To estimate the range of concentrations of the above-said 7 antibiotics that may define the sub-MIC selective window for *P. aeruginosa* PA14, we recorded growth curves of this bacterium in presence of 1/100, 1/50, 1/25, 1/10, 1/5 and 1/2 of its MICs to those antimicrobials (Figure 1) (see Methods). 1/200 of MIC was added in the case of ciprofloxacin, given the broad width of its selective window reported in other bacteria (Gullberg et al. 2011). Taking into account that MSC is related to the effect on fitness of the selective pressure (Hughes and Andersson 2012), the elected concentrations were the ones that reduced the fitness of *P. aeruginosa* PA14, either growth rate, final O.D. reached, or both traits; and also the concentration immediately below that did not modify those parameters, in order to screen a wider scope. In this way, we identified the selective conditions under which resistant mutants could more easily emerge and displace parental strain, based on their differential fitness in presence of the selective antibiotic. Data presented in Figure 1 support the hypothesis that quinolones, tetracycline and, to a lesser extent, amikacin sub-MIC selective window might be the widest, because a large number of concentrations severely hindered *P. aeruginosa* growth; as opposed to imipenem and polymyxin B, in which the concentrations range that modified bacterial fitness was the smallest. This analysis drove us to use 1/200 to 1/2 of MIC range for ciprofloxacin; 1/100 to 1/2 of MIC range for amikacin, levofloxacin and tetracycline; 1/50 to 1/2 of MIC range for ceftazidime; and 1/10 to 1/2 of MIC range for polymyxin B and imipenem; in the subsequent ALE.

**Figure 1.**
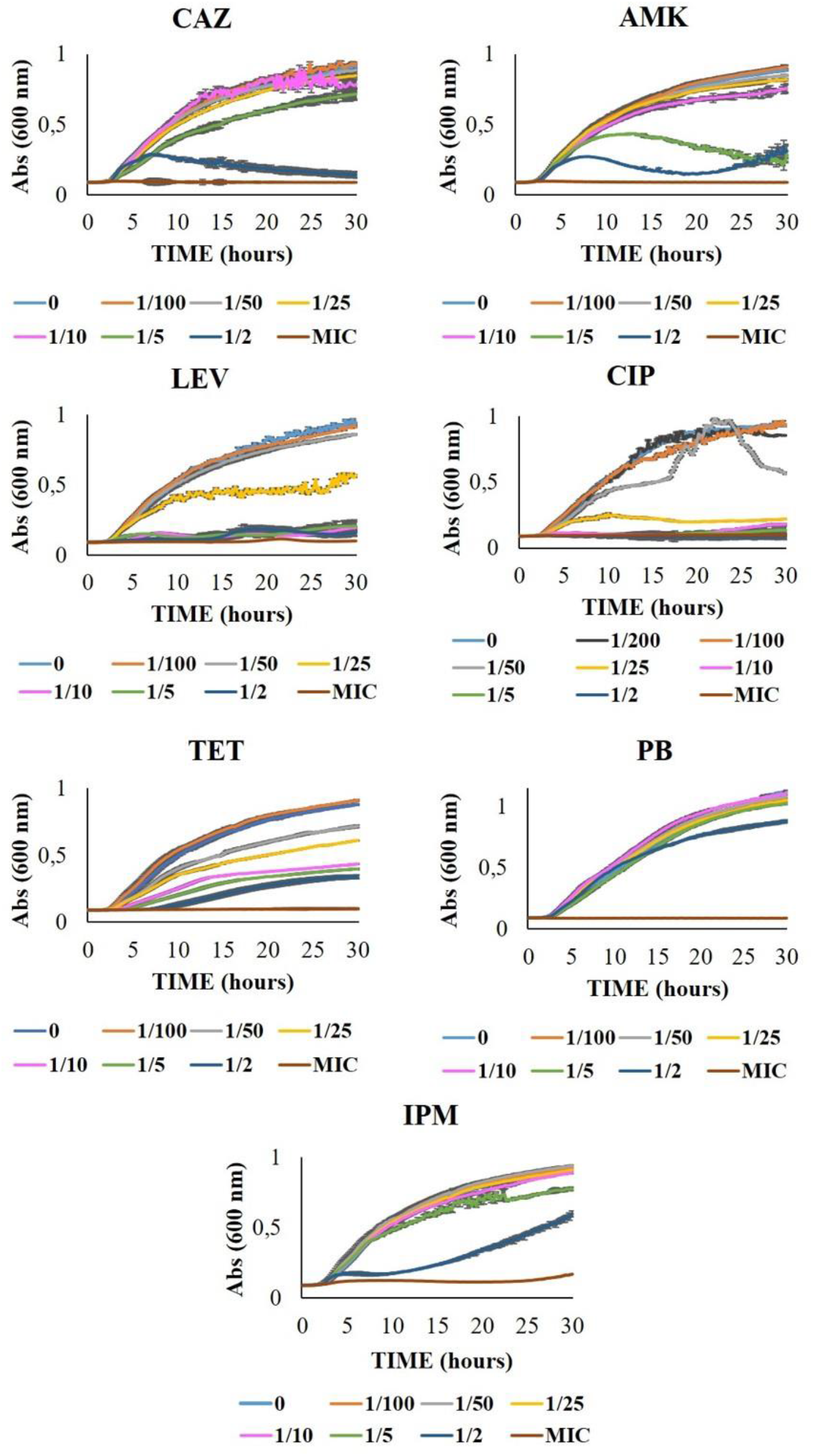
Growth curves of *P. aeruginosa* in presence of sub-MIC concentrations of 7 antibiotics. This graph shows the fitness of *P. aeruginosa* PA14 under 1/100, 1/50, 1/25, 1/10, 1/5 and 1/2 (plus 1/200 in the case of ciprofloxacin) of its MIC to 7 antibiotics. Experiments were performed in LB or MH II (in the case of polymyxin B) medium, for 30 hours at 37 °C. Error bars indicate standard deviations of the results from three independent replicates. CAZ: ceftazidime, AMK: amikacin, LEV: levofloxacin, CIP: ciprofloxacin, TET: tetracycline, PB: polymyxin B, IPM: imipenem.

### Sub-MIC selective windows of quinolones and amikacin are notably wide and select for clinically-relevant antibiotic resistant mutants

We undertook short-term ALE assays (9 days) of *P. aeruginosa* PA14, using seven different drugs, four replicates for each concentration and four control populations grown in absence of selective pressure, to shed light on the width of the sub-MIC selective window of each antimicrobial agent (Figure 2, Table S1). A sub-MIC concentration was considered as part of the selective window when, at least, half of the replicates evolved under those conditions became resistant (≥2-fold of the parental strain MIC value) at the end of the ALE. Although the majority of populations already acquired resistance in 3 days, some of them -notably the ones evolved in the presence of tetracycline or imipenem-lost that phenotype along the ALE. Given the instability of these AR phenotypes, we decided to establish 9 days of ALE as the time period to delimit the mutational-driven sub-MIC selective windows. In agreement with past findings (Ching and Zaman 2020), an effect on bacterial fitness caused by sub-MIC selective pressure did not always render AR selection at the end of the ALE, as we ascertained with 1/50 and 1/25 of MIC of tetracycline or 1/5 of MIC of polymyxin B, for instance. Howbeit, all concentrations that shaped the windows reduced as well PA14 fitness, as seen in the growth curves determined before evolution (Figure 1).

**Figure 2.**
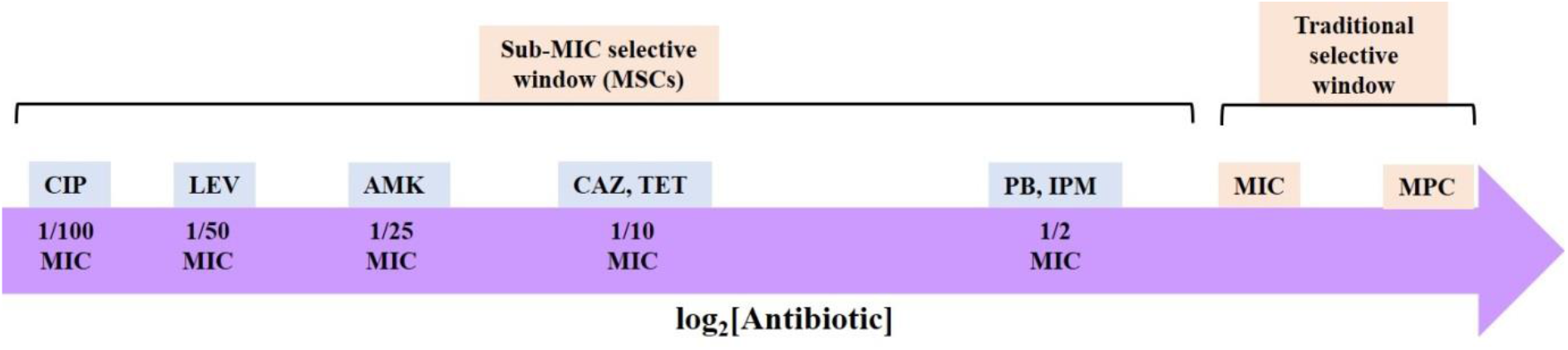
Schematic representation of the sub-MIC selective windows’ width for 7 antibiotics in *P. aeruginosa*. These data were ascertained by evolving *P. aeruginosa* PA14 populations for 9 days under a range of sub-MIC concentrations of each drug. A sub-MIC concentration was included in the selective window when at least half of the replicates evolved under those conditions became resistant (≥2-fold of the wild-type MIC value) at the end of the evolution period. MIC values for each replicate and controls are provided in Table S1. MSC: minimal selective concentration, MIC: minimal inhibitory concentration, MPC: mutant prevention concentration, CAZ: ceftazidime, AMK: amikacin, LEV: levofloxacin, CIP: ciprofloxacin, TET: tetracycline, PB: polymyxin B, IPM: imipenem.

The first idea distilled from the results presented in Figure 2 is that sub-MIC selective windows are antibiotic-specific, because 5 different widths were annotated, distributed among the 7 antimicrobials. As depicted there, quinolones present the widest window (1/100 and 1/50 of MIC select for resistant mutants to ciprofloxacin and levofloxacin, respectively), whilst polymyxin B and imipenem show the narrowest ones (1/2 of MIC is the only condition that allows AR selection). The large mutational space exhibited by the quinolones in this study sides with previous data about *Escherichia coli* sub-MIC selective window for ciprofloxacin (1/230 of the parental strain MIC) (Gullberg et al. 2011), and it is consistent with the broad range of concentrations of this drug that impaired *P. aeruginosa* fitness (Figure 1). Regarding the other antibiotics, ceftazidime and tetracycline sub-MIC selective windows reached 1/10 of MIC, whereas the one of amikacin spanned from 1/25 to 1/2 of MIC (Figure 2, Table S1). The latter is an interesting result, since we have recently established that tobramycin -a similar aminoglycoside-does not select *P. aeruginosa* PA14 resistant mutants in the presence of 1/10 of its MIC (Sanz-Garcia et al. 2020). This means that antibiotics belonging to the same structural family may present different sub-MIC selective windows and thus the risk of environmental pollution by one or another might differ. In the case here analysed, the environmental risk of amikacin (Tahrani et al. 2016), which presents a wider selective window, would be higher than tobramycin’s.

It is important to emphasize that quinolones were also the antibiotics that selected for the mutants exhibiting the highest increase in their resistance levels. To be precise, ciprofloxacin-evolved replicates typically reached ∼32-fold of the MIC of PA14 strain, up to 128-fold, and levofloxacin-evolved ones showed a resistance rise by regularly ∼21-fold, up to 168-fold (Table S1). Then again, imipenem and polymyxin B displayed, comparatively, a disparate tendency: the populations that became resistant to the first one only doubled their MICs with respect to the wild-type strain, whereas the two resistant replicates evolved under 1/2 of MIC of polymyxin B (P10, P11) reached 2-fold and 6-fold of the parental strain MIC, respectively. With regards to the remaining antibiotics, populations evolved under ceftazidime and amikacin sub-MIC selective windows increased their MICs up to 12-fold in both cases; and up to 8-fold in the case of tetracycline, respect to *P. aeruginosa* PA14 MIC.

Although in several cases (especially within ceftazidime, amikacin and ciprofloxacin’s selective windows) the levels of resistance correlated with the concentrations of selection, our results show that low antibiotic concentrations -even the lowest in the selective windows, i. e., the MSCs-may select clinically relevant antibiotic resistant mutants, presenting MICs above EUCAST clinical breakpoints. This situation was observed in levofloxacin, ciprofloxacin, amikacin and, to a lesser extent, ceftazidime ALEs. In this regard, the study of quinolones provided, once again, the most remarkable result: 1/50 of MIC, a concentration that can be encountered in non-clinical habitats (see below), was enough pressure to exert selection for clinically resistant mutants to ciprofloxacin and levofloxacin (Table S1). Hence, it could be generally asserted that low concentrations of antibiotics -as long as they belong to the selective window-may select for highly resistant mutants of *P. aeruginosa* with clinical relevance, according to EUCAST standards.

### Sub-MIC concentrations of all tested drugs select for cross-resistance to antibiotics of clinical value

We have formerly reported that *P. aeruginosa* resistant mutants selected under sub-MIC tigecycline concentrations (down to 1/50 of PA14 MIC) exhibit cross-resistance to other antibiotics besides the one used along selection (Sanz-Garcia et al. 2020). In order to decipher whether this phenomenon also occurs under the sub-MIC selective windows of the 7 drugs used in this work, the MICs of a set of antibiotics from distinct structural families were determined for the final evolved populations which resistance level to the selective antibiotic was ≥2-fold of the parental strain MIC value.

Every resistant replicate evolved under their designated sub-MIC selective window showed lower susceptibility to at least one antimicrobial different from the selective one, especially to quinolones, tetracycline, chloramphenicol and erythromycin (Figure 3, Table S2). In fact, exposure to all selective antibiotics led to up to 7 cases of cross-resistance to antibiotics belonging to dissimilar families, some of which are of crucial value in the treatment of *P. aeruginosa* (i. e. ceftazidime, aztreonam, amikacin or levofloxacin) (Bassetti et al. 2018). Overall, quinolones selected for fewer cases of cross-resistance, suggesting that they may have selected more specific mechanisms of resistance than the ones selected by the other drugs analyzed in the current study.

**Figure 3.**
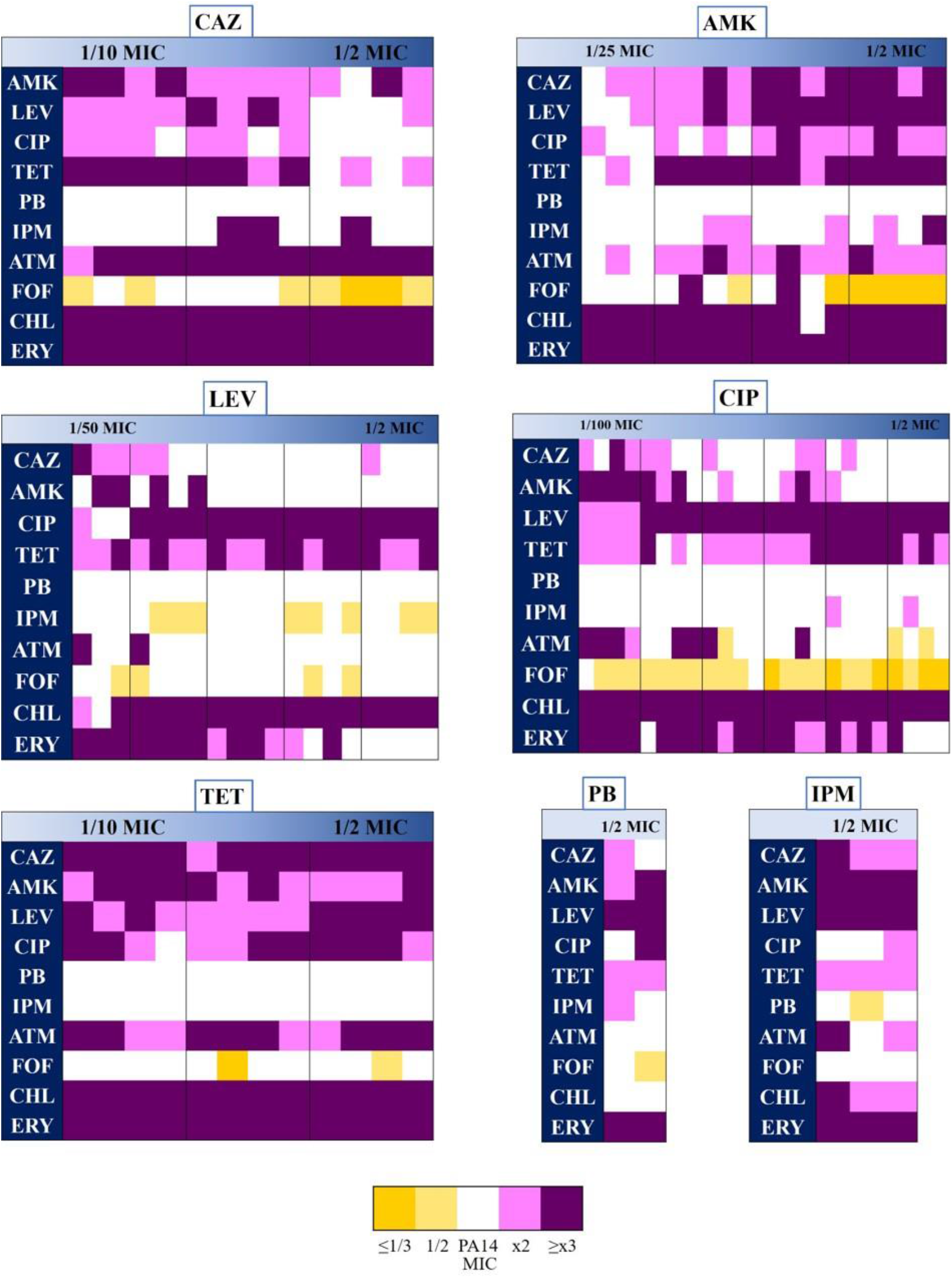
Cross-resistance and collateral sensitivity phenotypes of populations evolved under sub-MIC selective windows of 7 antibiotics. Cross-resistance and collateral sensitivity to antibiotics from different structural families were analysed in the populations that had become resistant (≥2-fold of the wild-type MIC value) at the end of the ALEs. A population was labelled as susceptible (yellow) or resistant (purple) when there was a MIC change, at least by 1/2 or 2-fold, with respect to the wild-type strain value. MIC values are encompassed in Table S2. CAZ: ceftazidime, AMK: amikacin, LEV: levofloxacin, CIP: ciprofloxacin, TET: tetracycline, PB: polymyxin B, IPM: imipenem, ATM: aztreonam, FOF: fosfomycin, CHL: chloramphenicol, ERY: erythromycin.

In certain cases, the strength of these cross-resistance phenotypes positively correlated with the concentration of the selective drug (i. e., higher sub-MIC concentrations of amikacin selected for higher cross-resistance to quinolones), albeit in many others there was not such correlation (i. e., tetracycline sub-MIC ALE assays led to similar levels of ceftazidime and erythromycin cross-resistance in the entire sub-MIC selective window) (Table S2). These results indicate that there is not a general quantitative correlation between the level of resistance to the selective antibiotic and the level of cross-resistance to antimicrobial agents belonging to different categories, a feature in line with previous findings (Sanz-Garcia et al. 2020). To note here that none of the final evolved populations presented polymyxin B cross-resistance, and only two displayed resistance to fosfomycin (Figure 3). As a matter of fact, some sub-MIC concentrations of ceftazidime, amikacin and especially ciprofloxacin, selected for collateral sensitivity to fosfomycin, whereas levofloxacin sub-MIC selective window drove to several cases of slight collateral sensitivity to imipenem. Hyper-susceptibility to fosfomycin in *P. aeruginosa* PA14 has been described in former ALE studies in our laboratory, precisely in presence of ceftazidime and the aminoglycoside tobramycin, among other drugs (Hernando-Amado et al. 2020; Sanz-Garcia et al. 2018a, b), likely due to reduction in the expression of *fosA* -which codes for an enzyme that degrades fosfomycin- and genes encoding enzymes involved in the peptidoglycan recycling pathway (Laborda et al. 2021, submitted).

We ought to remark that not only concentrations of these antibiotics as low as 1/100 of MIC (i. e., ciprofloxacin) may select for MDR *P. aeruginosa* mutants, but even some of these concentrations could select for particularly high levels of cross-resistance that equate or surpass the EUCAST clinical breakpoints for this pathogen. This is the case of aztreonam or imipenem MICs of certain ceftazidime-evolved resistant populations, or levofloxacin in ciprofloxacin-evolved resistant ones -and vice versa-(Table S2).

Altogether, these data hint that antibiotic concentrations encompassed in the sub-MIC selective window are capable of giving rise to mutants presenting joint resistance to different drugs that are essential in antipseudomonal therapies.

### MDR patterns in mutants selected under sub-MIC selective windows may be prevalently caused by efflux pumps’ activity

Antibiotic cross-resistance is frequently due to mutations that increase or modify the activity of MDR efflux pumps (Blair et al. 2015; Blanco et al. 2019; Hernando-Amado et al. 2016). To determine whether these systems could have a role in the observed MDR phenotypes, the MICs for the resistant populations to some specific antimicrobials were elucidated in presence of the EPI PAβN (Lomovskaya et al. 2001), and compared with the ones obtained without it. PAβN is a non-specific inhibitor of RND-type efflux systems, and its effect on *P. aeruginosa*’s susceptibility to almost every drug tested in this work has been extensively studied (Kao et al. 2016; Mao et al. 2001; Renau et al. 2002; Sonnet et al. 2012; Xu et al. 2014). We chose the antibiotics based on the following criteria: selective antibiotic must be one of them, they must belong to different families, and at least half of the replicates evolved under each sub-MIC concentration must show resistance (≥2-fold of the wild-type MIC value) against them. Also, we included the drugs that fitted the substrate range of the most relevant *P. aeruginosa* efflux pumps, since mutations leading to their overexpression may have been selected in the presence of each sub-MIC ALE. We considered that resistance was at least partly efflux-mediated when MIC in presence of PAβN was reduced to ≤1/2 of the value in absence of that compound. Albeit, it must be noted that parental strain’s MICs to levofloxacin, chloramphenicol and erythromycin already decreased in presence of PAβN (Table S3), presumably because of the inhibition of MexAB-OprM and/or MexXY-OprN activity, which contribute to *P. aeruginosa*’s intrinsic resistance to these drugs (Masuda et al. 2000a; Olivares et al. 2013). In such cases, resistance of an evolved population was labelled as “efflux-mediated” when PAβN reduced its MIC to the PA14 level in presence of the EPI as well.

The results of this experiment support that MDR efflux pumps have a prevailing involvement in the AR phenotype selected by sub-MIC concentrations of several antibiotics in *P. aeruginosa*. As shown in Figure 4 and Table S3, efflux pumps’ activity appears to hold a prominent role in most cases of resistance to ceftazidime, ciprofloxacin, tetracycline and imipenem, which buttresses the likely importance of these systems in selection of *de novo* resistant mutants under low concentrations of drugs. Some populations showed reduced MICs in presence of PAβN but still maintained a higher value (≥2-fold) than the ones of PA14. Here, other resistance mechanisms, in addition to efflux pumps, might be contributing to AR.

**Figure 4.**
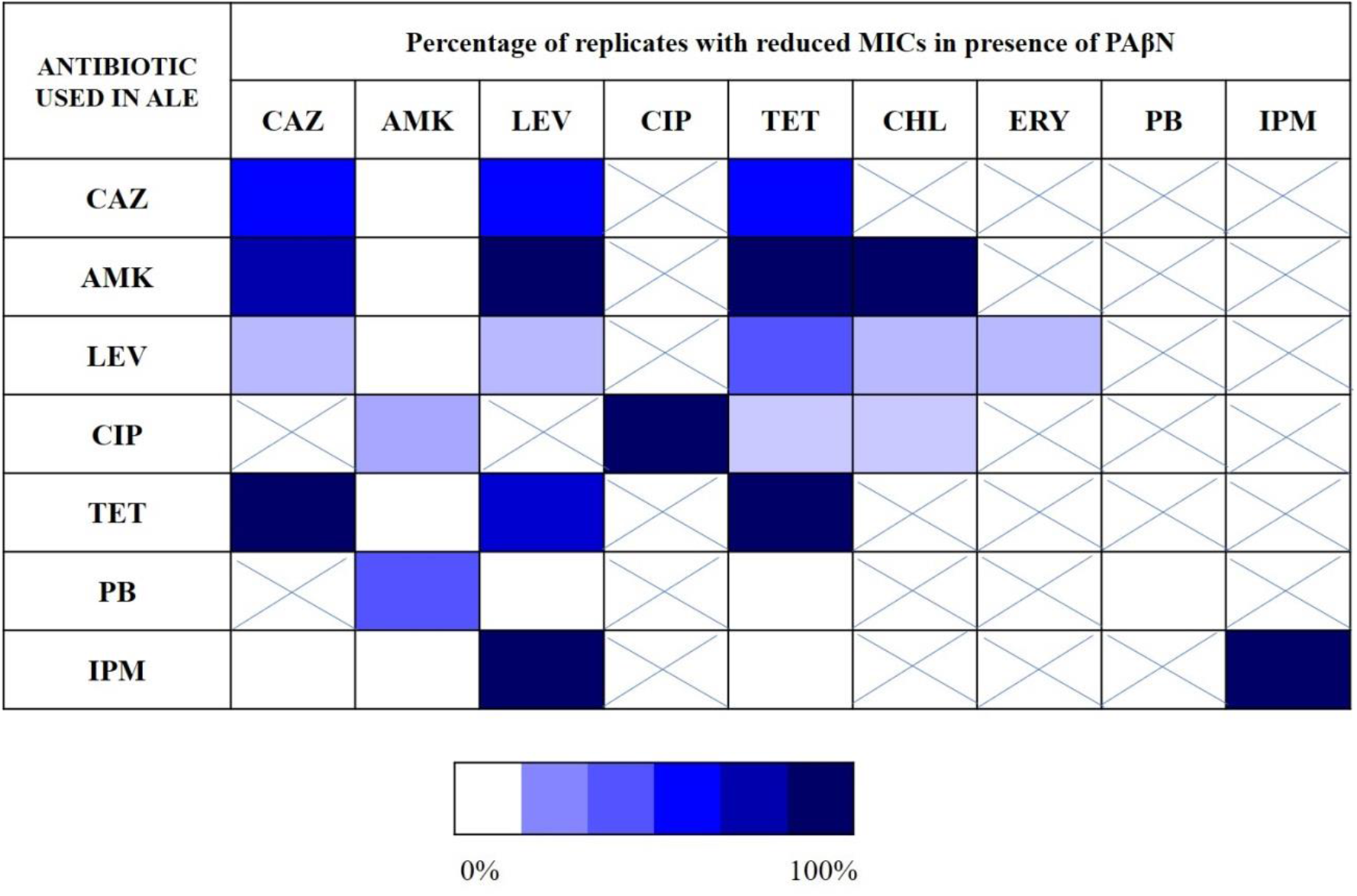
Role of MDR efflux pumps in AR phenotypes of populations evolved under sub-MIC selective windows of 7 antibiotics. MICs to different antibiotics in presence of the EPI PAβN were analysed in the populations that had become MDR at the end of the ALEs, and compared with the MIC values in absence of EPI. Blue intensity code indicates the percentage of replicates evolved under each drug that showed a MIC decrease in presence of PAβN (≤1/2), when compared to the value without EPI. An “X” indicates the antimicrobials whose MICs were not measured in this analysis. MIC values are encompassed in Table S3. CAZ: ceftazidime, AMK: amikacin, LEV: levofloxacin, CIP: ciprofloxacin, TET: tetracycline, CHL: chloramphenicol, ERY: erythromycin, PB: polymyxin B, IPM: imipenem.

It should be mentioned that all populations resultant from 1/2 of MIC of ceftazidime ALE were unable to grow in presence of EPI, impeding us to know the contribution of efflux pumps to their resistance patterns (Table S3). We hypothesize that, since 3 out of 4 (C18-C20) of these populations hyper-produced the dark-brown pigment pyomelanin, which is a visual marker of large chromosomal deletions selected by ceftazidime in *P. aeruginosa* PA14 (Sanz-Garcia et al. 2018a), the lack of some genes may have provoked an impossibility to thrive in presence of this inhibitor of efflux activity, given its broad effect on bacterial physiology (Rampioni et al. 2017). However, this is a feature that remains to be further analyzed. These chromosomal deletions usually provide with β-lactams resistance (Alvarez-Ortega et al. 2010) that may be associated with an increased β-lactamase activity. Therefore, we measured β-lactamase activity in the C17-C20 population replicates. All the populations showed a statistically significant enhancement of β-lactamase activity with respect to the wild-type strain (Figure 5), which may explain their resistance to ceftazidime and aztreonam. Nevertheless, unraveling the exact reason behind cross-resistance phenotypes of these populations will demand further investigation.

**Figure 5.**
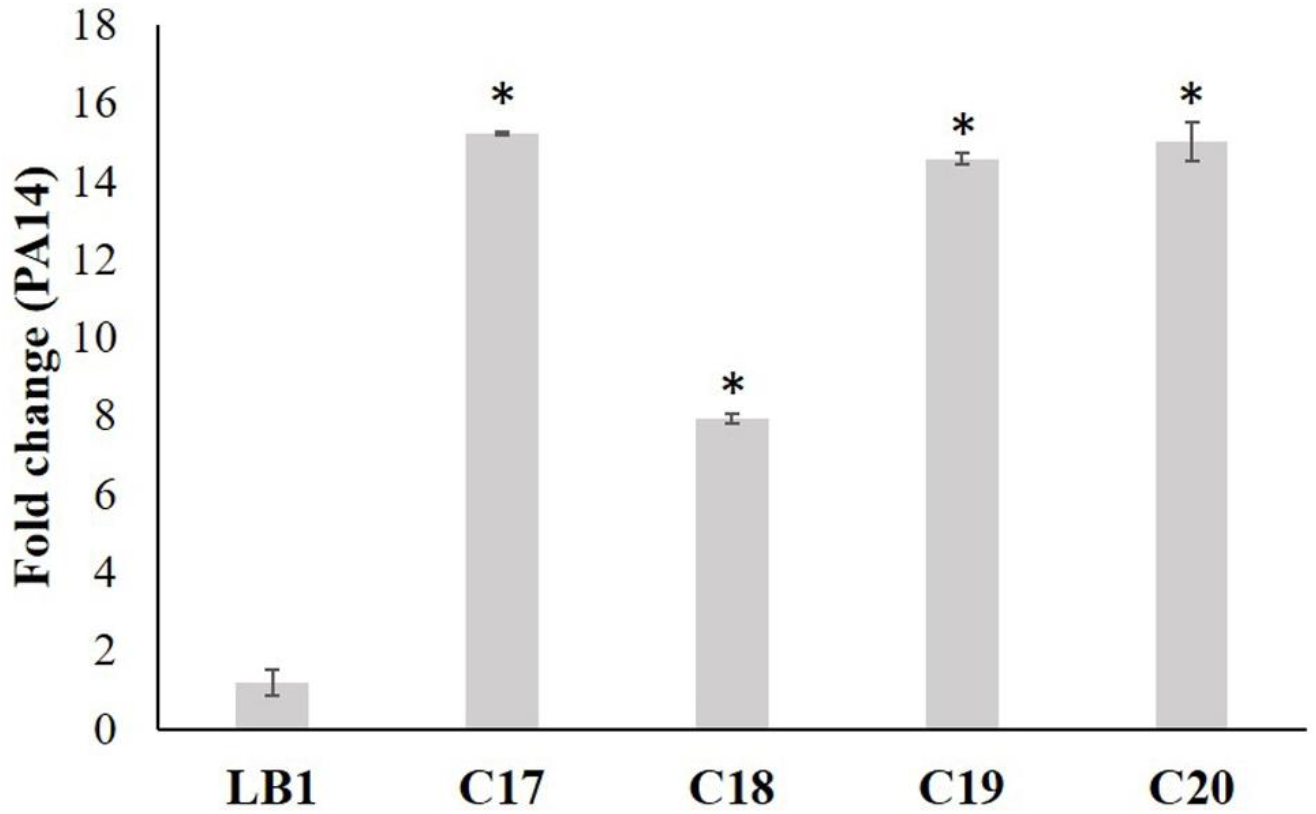
β-lactamase activity in populations evolved under sub-MIC selective windows of 7 antibiotics. Relative amount of β-lactamase activity in populations evolved under 1/2 of MIC of ceftazidime (C17-C20) and a control replicate evolved in absence of antibiotic (LB1). Fold changes were estimated with respect to the value of *P. aeruginosa* PA14 strain. Error bars indicate standard deviations of the results from two independent experiments. Statistically significant differences (*P* < 0,05) were evaluated using Student’s *t-*test and are highlighted with an asterisk (*).

Apropos of levofloxacin and polymyxin B-evolved resistant populations, the effect of the EPI on their MICs was minor, suggesting a marginal role of efflux in their AR phenotypes. In regard to the first ones, only the replicates submitted to 1/50 of MIC of levofloxacin showed a consistent increase in susceptibility in presence of PAβN. Tetracycline cross-resistance of a levofloxacin-resistant population evolved in the presence of 1/5 of MIC and of all the populations evolved in the presence of 1/2 of MIC was also impaired by the EPI. Concerning polymyxin B, only one replicate (out of 2) evolved under 1/2 of MIC of this polycationic compound showed a diminishment of amikacin resistance when PAβN was present. We could conjecture this may be a consequence of PAβN effect on membrane permeability (Lamers et al. 2013), which could explain amikacin MIC reduction.

## DISCUSSION

Low antibiotic concentrations found in nature, released from anthropogenic sources - beside the ones in clinical settings-, are a pervasive and often neglected threat to human health, because they could lead to selection of highly resistant bacteria (Wistrand-Yuen et al. 2018) if they fit within their sub-MIC selective windows. Despite *P. aeruginosa* being one of the most ubiquitous pathogens in natural environments, works have customarily focused on delineating its traditional above-MIC mutant selective window to antimicrobial agents (Kawamura et al. 2019; Morero et al. 2011; Xu et al. 2014; Zinner et al. 2013), whilst MSC determination has only been performed in other bacteria and just for a reduced number of drugs (Kraupner et al. 2018; Kraupner et al. 2020; Lundstrom et al. 2016; Murray et al. 2018; Stanton et al. 2020). Studies about the effect of ciprofloxacin sub-MIC concentrations have been previously undertaken in *P. aeruginosa*, but most of them used higher concentrations than the ones here applied (Jorgensen et al. 2013). Consequently, as far as we know, this is the first work to unveil the minimal sub-MIC concentration of clinically valuable antibiotics (most of them belonging to regular antipseudomonal therapies) that are able to select for *P. aeruginosa* resistant mutants (Table 1).

**Table 1.** Overview of the MSC values (µg/l) of *P. aeruginosa* to 7 antibiotics, their presence in nature and their effect on this pathogen.

| Antibiotic | MSC (µg/l) | Highest concentration found in nature (µg/l) (ref) | MDR phenotype | Implication of efflux pumps in MDR phenotype |
| --- | --- | --- | --- | --- |
| CAZ | 400 | ND | AMK, LEV, CIP, TET, ATM, CHL, ERY | Yes |
| AMK | 160 | 2.3<br>(Tahrani et al. 2016) | CHL, ERY | Yes |
| LEV* | 80 | 196<br>(Mohapatra et al. 2016) | CAZ, AMK, TET, CHL, ERY | Yes |
| CIP* | 5 | 31000<br>(Larsson et al. 2007) | CAZ, AMK, LEV, TET, ATM, CHL, ERY | Yes |
| TET* | 1500 | 60790<br>(Hou et al. 2016) | CAZ, AMK, LEV, CIP, ATM, CHL, ERY | Yes |
| PB | 2000 | ND | CAZ, AMK, LEV, CIP, TET, IPM, ERY | No |
| IPM | 250 | ND | CAZ, AMK, LEV, TET, ATM, CHL, ERY | Yes |

The table encloses the MSC values of *P. aeruginosa* to 7 drugs, the highest amount of them that has been detected in nature, the MDR this concentration selects for (at least half the replicates must show $\geq 2$ -fold of the parental strain MIC to be subsumed), and the potential implication of MDR efflux pumps in contributing to this phenotype. \*: MSC of this antibiotic is lower than its highest concentration found in nature. ND: No data. CAZ: ceftazidime, AMK: amikacin, LEV: levofloxacin, CIP: ciprofloxacin, TET: tetracycline, PB: polymyxin B, IPM: imipenem, ATM: aztreonam, CHL: chloramphenicol, ERY: erythromycin.

In agreement with previous findings (Gullberg et al. 2011; Sanz-Garcia et al. 2020), our results show that sub-MIC selective windows are antibiotic-specific, being the ones of quinolones the largest, and the ones of imipenem and polymyxin B the narrowest. In addition, our results show that many of the extremely low concentrations of drugs tested enrich bacterial populations with clinically relevant MDR mutants, presenting MICs beyond clinical breakpoints (Tables S1-S2). In spite of being previously discussed that low-level resistant mutants should predominate at sub-MIC antibiotic conditions (Martinez and Baquero 2000), studies based in *de novo* selection of resistant mutants under different drug concentrations (Garcia-Leon et al. 2014) and in the analysis of the fitness of reconstructed resistant mutants growing on sub-MIC antibiotic concentrations (Gullberg et al. 2011), have proved that clinically relevant AR may be selected even at low antimicrobial concentrations. Our findings concur with these studies, at least for the analyzed antibiotics.

From a One-Health perspective, we should underline that, among the antibiotics chosen for this study, there is information about ciprofloxacin, levofloxacin, tetracycline and amikacin presence in aquatic ecosystems in which *P. aeruginosa* may thrive (Danner et al. 2019; Hou et al. 2016; Mena and Gerba 2009; Mohapatra et al. 2016; Rodriguez-Mozaz et al. 2020; Tahrani et al. 2016). From this set, ciprofloxacin stands out, considering that high concentrations of this antibiotic have been detected in many ecosystems: 13 µg/l or 27 µg/l, up to 6.5 mg/l or 31 mg/l of this compound have been reported to pollute Spain hospital effluents (Rodriguez-Mozaz et al. 2015), South Africa stream waters (Agunbiade and Moodley 2016), a lake from India (Fick et al. 2009), and a pharmaceutical plant effluent in the same country (Larsson et al. 2007), respectively. In the current work, we observe that 5 µg/l may already select for *P. aeruginosa* resistant mutants to ciprofloxacin, amikacin or aztreonam (Table 1). Actually, 10 µg/l of ciprofloxacin already fosters the emergence of clinically resistant mutants to this quinolone, following EUCAST criteria. Moreover, according to our experiments, the MSCs of levofloxacin and tetracycline (80 µg/l and 1.5 mg/l), are lower than specific amounts of these drugs that have been measured in wastewater treatment plants in India (196 µg/l) (Mohapatra et al. 2016) and Northern China (2.6, 11 or 60.79 mg/l) (Hou et al. 2016), respectively (Table 1). To note here that *P. aeruginosa* strains have been found to dominate pharmaceutical production environments (Ratajczak et al. 2021), which drug pollution levels are usually the highest. Thus, the information comprised in this work concerning quinolones effect on this opportunistic pathogen is considerably worrying -especially respecting ciprofloxacin-, given their wide sub-MIC selective window, the elevated levels of resistance to the selective antibiotic and cross-resistance to other drugs that selected mutants display, and the certified pollution by these drugs in several habitats, in concentrations that fit within the selective windows determined in the current work. Regarding tetracycline, in spite of not being used against *P. aeruginosa* infections, it is disquieting that its concentration in aquatic habitats might be able to select for resistant mutants to drugs of clinical value, as ceftazidime or aztreonam (Table 1). This result exemplifies the possibility of selecting resistance to antibiotics of clinical relevance even in environments not polluted by them. Besides these non-clinical situations, it should not be disregarded that, in certain regions, poor-quality medicines, which harbour a substandard quantity of API, are being used in clinics, being precisely the case of ciprofloxacin and tetracycline (Frimpong et al. 2018; Johnston and Holt 2014; Tabernero et al. 2019). Then, this reality enlarges the variety of circumstances under which AR selection mediated by low antimicrobial concentrations could be befalling.

Although, to the best of our knowledge, neither information about the amount of ceftazidime, polymyxin B and imipenem present in natural environments (Umweltbundesamt 2016), nor the usage of substandard medicines containing them, have been documented, the fact that all antibiotics used for human therapy or farming are regularly released in water and soils, supports these antibiotics to be present in habitats frequently colonized by *P. aeruginosa* (Morales et al. 2004). Nonetheless, the narrowness of imipenem and polymyxin B sub-MIC selective windows (Figure 2) suggests that pollution by these drugs would be less dangerous from a One-Health view, because their sub-MIC mutational space is eminently reduced compared to the one of the aforementioned quinolones, so that the chances of selecting resistant mutants in clinical and non-clinical settings harbouring these drugs are lower.

With the aim of addressing the implication of MDR efflux pumps in AR selected under sub-MIC antibiotic concentrations, we used an EPI (PAβN). As depicted in Figure 4, and as one should expect in light of so many cross-resistances found, efflux activity seems to play a key role in the observed MDR phenotypes, across most drugs and concentrations ranges. This result sides with prior tetracycline and ciprofloxacin sub-MIC assays, in which MexXY-OprM, MexAB-OprM and/or MexCD-OprJ were found overexpressed (C. Barbosa et al. 2021; Morero et al. 2011; Morita et al. 2006). To note here that the substrate range of the efflux pumps these mutants might overexpress, explains the nearly total absence of polymyxin B and fosfomycin cross-resistance across the evolved replicates, since these compounds are not usual substrates of most *P. aeruginosa*’s pumps. In addition, it should be commented that MDR patterns which, according to the EPI experiment performed, were not caused by efflux systems (the ones of most levofloxacin-evolved replicates and one polymyxin B-evolved replicate), could be the consequence of mutations that altered membrane permeability, which might trigger pleiotropic effects.

In conclusion, the results here encompassed allow us to predict the level of resistance, cross-resistance and collateral sensitivity of *P. aeruginosa* mutants that might be selected for under specific low concentrations of 7 different drugs that can be encountered in natural (non-clinical) ecosystems. This information could constitute a useful asset in the fight against AR from a One-Health and Global-Health perspective.

## Acknowledgements

Work in the laboratory is supported by grant S2017/BMD-3691 InGEMICS-CM, funded by Comunidad de Madrid (Spain) and European Structural and Investment Funds, by Instituto de Salud Carlos III (grant RD16/0016/0011) - cofinanced by the European Development Regional Fund “A Way to Achieve Europe”, and by the Spanish Ministry of Economy and Competitivity (BIO2017-83128-R). The funders had no role in study design, data collection and interpretation, or the decision to submit the current work for publication. The authors declare that they have no conflict of interest.

