## Supplemental Data for "Evolution under low antibiotic concentrations: a risk for the selection of *Pseudomonas aeruginosa* multidrug resistant mutants in nature"

**Table S1 | MIC values (µg/ml) of *P. aeruginosa* PA14 populations during short-term ALEs under sub-MIC concentrations of 7 different antibiotics.**

| | | | TIME (days) | | | |
|---|---|---|---|---|---|---|
| **ANTIBIOTIC** | **CONCENTRATION (relative to MIC)** | **REPLICATE** | **0** | **3** | **6** | **9** |
| **CAZ (8)** | **1/50** | **C1** | 0,75 | 1 | 1 | 0,75 |
| | | **C2** | 0,75 | 0,75 | 1 | 1 |
| | | **C3** | 0,75 | 1 | 1 | 0,75 |
| | | **C4** | 0,75 | 1 | 1 | 1 |
| | **1/25** | **C5** | 0,75 | 1 | 1 | 1 |
| | | **C6** | 0,75 | 1 | 1 | 1 |
| | | **C7** | 0,75 | 0,75 | 0,75 | 1 |
| | | **C8** | 0,75 | 1 | 1 | 1 |
| | **1/10** | **C9** | 0,75 | 3 | 3 | 2 |
| | | **C10** | 0,75 | 3 | 2 | 2 |
| | | **C11** | 0,75 | 3 | 2 | 2 |
| | | **C12** | 0,75 | 3 | 3 | 3 |
| | **1/5** | **C13** | 0,75 | 3 | 6 | 4 |
| | | **C14** | 0,75 | 3 | 3 | 2 |
| | | **C15** | 0,75 | 4 | 6 | 8 |
| | | **C16** | 0,75 | 3 | 6 | 4 |
| | **1/2** | **C17** | 0,75 | 4 | 8 | 6 |
| | | **C18** | 0,75 | 4 | 12 | 12 |
| | | **C19** | 0,75 | 4 | 6 | 8 |
| | | **C20** | 0,75 | 3 | 4 | 6 |
| **AMK (16)** | **1/100** | **A1** | 2 | 4 | 4 | 3 |
| | | **A2** | 2 | 3 | 3 | 3 |
| | | **A3** | 2 | 3 | 2 | 2 |
| | | **A4** | 2 | 3 | 3 | 2 |
| | **1/50** | **A5** | 2 | 3 | 3 | 3 |
| | | **A6** | 2 | 2 | 2 | 3 |
| | | **A7** | 2 | 3 | 2 | 3 |
| | | **A8** | 2 | 2 | 2 | 3 |
| | **1/25** | **A9** | 2 | 3 | 4 | 4 |
| | | **A10** | 2 | 2 | 3 | 2/4 |
| | | **A11** | 2 | 3 | 3 | 3 |
| | | **A12** | 2 | 4 | 4 | 3/6 |
| | **1/10** | **A13** | 2 | 3/8 | 2/6 | 6 |
| | | **A14** | 2 | 2/4 | 4/8 | 3/8 |
| | | **A15** | 2 | 3/8 | 12 | 16 |
| | | **A16** | 2 | 2/6 | 2/6 | 12 |

| | | | | | | |
|---|---|---|---|---|---|---|
| | **1/5** | **A17** | 2 | 8 | 6/16 | 6/24 |
| | | **A18** | 2 | 16 | 6/16 | 16 |
| | | **A19** | 2 | 8 | 12 | 8 |
| | | **A20** | 2 | 12 | 12/24 | 24 |
| | **1/2** | **A21** | 2 | 8/32 | 32 | 24 |
| | | **A22** | 2 | 16/64 | 48 | 24 |
| | | **A23** | 2 | 48 | 48 | 24 |
| | | **A24** | 2 | 64 | 32 | 24 |
| **LEV (1)** | **1/100** | **L1** | 0,19 | 0,19 | 0,25 | 0,25 |
| | | **L2** | 0,19 | 0,19 | 0,25 | 0,25 |
| | | **L3** | 0,19 | 0,19 | 0,38 | 0,25 |
| | | **L4** | 0,19 | 0,19 | 0,25 | 0,25 |
| | **1/50** | **L5** | 0,19 | 0,19/1,5 | 0,19 | 0,25/0,5 |
| | | **L6** | 0,19 | 0,094/1 | 0,19 | 0,19/0,5 |
| | | **L7** | 0,19 | 0,19/1,5 | 0,19/1,5 | 0,19/2 |
| | | **L8** | 0,19 | 0,19 | 0,19 | 0,25 |
| | **1/25** | **L9** | 0,19 | 4 | 3 | 2 |
| | | **L10** | 0,19 | 4 | 3 | 0,75/3 |
| | | **L11** | 0,19 | 4 | 3 | 3 |
| | | **L12** | 0,19 | 4 | 4 | 4 |
| | **1/10** | **L13** | 0,19 | 6 | 3 | 3 |
| | | **L14** | 0,19 | 6 | 4 | 3 |
| | | **L15** | 0,19 | 6 | 4 | 3 |
| | | **L16** | 0,19 | 6 | 4 | 4 |
| | **1/5** | **L17** | 0,19 | 4 | 4 | 3 |
| | | **L18** | 0,19 | 4 | 3 | 4 |
| | | **L19** | 0,19 | 4 | 4 | 3 |
| | | **L20** | 0,19 | 4 | 4 | 4 |
| | **1/2** | **L21** | 0,19 | 4 | 8 | 8 |
| | | **L22** | 0,19 | 4 | 4 | 6 |
| | | **L23** | 0,19 | 4 | 8 | 12 |
| | | **L24** | 0,19 | 4/8 | ≥32 | ≥32 |
| | **1/200** | **X1** | 0,047 | 0,064 | 0,094 | 0,064 |
| | | **X2** | 0,047 | 0,094 | 0,064 | 0,064 |
| | | **X3** | 0,047 | 0,064 | 0,064 | 0,064 |
| | | **X4** | 0,047 | 0,094 | 0,064 | 0,094 |
| | **1/100** | **X5** | 0,047 | 0,064/1 | 0,094 | 0,125 |
| | | **X6** | 0,047 | 0,064 | 0,064 | 0,125 |
| | | **X7** | 0,047 | 0,064/0,5 | 0,094 | 0,094 |
| | | **X8** | 0,047 | 0,047/0,38 | 0,064 | 0,094 |
| | **1/50** | **X9** | 0,047 | 0,75 | 0,75 | 0,75 |
| | | **X10** | 0,047 | 1 | 0,75 | 0,75 |
| | | **X11** | 0,047 | 0,75 | 0,5 | 0,75 |

| | | | | | | |
|---|---|---|---|---|---|---|
| **CIP (0,5)** | | **X12** | 0,047 | 1 | 0,5 | 0,75 |
| | **1/25** | **X13** | 0,047 | 1,5 | 1 | 1 |
| | | **X14** | 0,047 | 1,5 | 1 | 1 |
| | | **X15** | 0,047 | 1 | 0,75 | 0,75 |
| | | **X16** | 0,047 | 1 | 1,5 | 1,5 |
| | **1/10** | **X17** | 0,047 | 1,5 | 1 | 1,5 |
| | | **X18** | 0,047 | 2 | 1 | 1,5 |
| | | **X19** | 0,047 | 1,5 | 1 | 1 |
| | | **X20** | 0,047 | 1 | 1 | 1,5 |
| | **1/5** | **X21** | 0,047 | 1 | 1 | 16 |
| | | **X22** | 0,047 | 1,5 | 1 | 3 |
| | | **X23** | 0,047 | 2 | 1,5 | 3 |
| | | **X24** | 0,047 | 2 | 1 | 4 |
| | **1/2** | **X25** | 0,047 | 1,5 | 2 | ≥32 |
| | | **X26** | 0,047 | 1,5 | 2 | 6 |
| | | **X27** | 0,047 | 2 | 2 | 12 |
| | | **X28** | 0,047 | 1,5 | 6 | 6 |
| **TET** | **1/100** | **T1** | 6 | 8 | 12 | 4 |
| | | **T2** | 6 | 12 | 8 | 6 |
| | | **T3** | 6 | 4 | 6 | 4 |
| | | **T4** | 6 | 12 | 8 | 4 |
| | **1/50** | **T5** | 6 | 12 | 6 | 6 |
| | | **T6** | 6 | 12 | 6 | 6 |
| | | **T7** | 6 | 12 | 8 | 6 |
| | | **T8** | 6 | 24 | 24 | 4 |
| | **1/25** | **T9** | 6 | 8 | 12 | 4 |
| | | **T10** | 6 | 16 | 8 | 6 |
| | | **T11** | 6 | 16 | 6 | 6 |
| | | **T12** | 6 | 12 | 8 | 6 |
| | **1/10** | **T13** | 6 | 16 | 16 | 12 |
| | | **T14** | 6 | 16 | 16 | 16 |
| | | **T15** | 6 | 16 | 24 | 24 |
| | | **T16** | 6 | 16 | 16 | 16 |
| | **1/5** | **T17** | 6 | 8 | 24 | 24 |
| | | **T18** | 6 | 12 | 24 | 24 |
| | | **T19** | 6 | 12 | 32 | 16 |
| | | **T20** | 6 | 16 | 32 | 24 |
| | **1/2** | **T21** | 6 | 24 | 24 | 24 |
| | | **T22** | 6 | 32 | 64 | 48 |
| | | **T23** | 6 | 16 | 24 | 24 |
| | | **T24** | 6 | 16 | 24 | 24 |
| | **1/10** | **P1** | 1 | 1,5 | 1 | 1 |
| | | **P2** | 1 | 1 | 0,75 | 1 |

| | | | | | | |
|---|---|---|---|---|---|---|
| **PB*** | | **P3** | 1 | 1 | 1 | 1,5 |
| | | **P4** | 1 | 1 | 1 | 1 |
| | **1/5** | **P5** | 1 | 1 | 1 | 1 |
| | | **P6** | 1 | 1 | 1 | 1 |
| | | **P7** | 1 | 1 | 1 | 1 |
| | | **P8** | 1 | 1,5 | 1 | 1 |
| | **1/2** | **P9** | 1 | 1,5 | 3 | 1,5 |
| | | **P10** | 1 | 1,5 | 2 | 2 |
| | | **P11** | 1 | 3/6 | 2/6 | 6 |
| | | **P12** | 1 | 1,5 | 1,5 | 1,5 |
| **IPM (4)** | **1/10** | **I1** | 1 | 1 | 0,75 | 0,5 |
| | | **I2** | 1 | 2 | 1 | 0,75 |
| | | **I3** | 1 | 2 | 1,5 | 0,75 |
| | | **I4** | 1 | 2 | 1,5 | 1 |
| | **1/5** | **I5** | 1 | 2 | 1,5 | 1 |
| | | **I6** | 1 | 2 | 1 | 1 |
| | | **I7** | 1 | 1,5 | 1 | 1 |
| | | **I8** | 1 | 2 | 1 | 1 |
| | **1/2** | **I9** | 1 | 1,5 | 2 | 2 |
| | | **I10** | 1 | 1,5 | 1,5 | 2 |
| | | **I11** | 1 | 1 | 1,5 | 2 |
| | | **I12** | 1 | 1 | 1,5 | 1 |

The table shows the MIC values, every 3 days, for each replicate population evolved under sub-MIC concentrations of 7 different antibiotics. Relative sub-MIC concentrations are determined based on MICs of *P. aeruginosa* PA14 in LB medium (or MH II, in polymyxin B's case). The concentrations belonging to the sub-MIC selective window are represented as a black cell. All MICs are obtained by E-test strips. Double inhibition halos are shown as two MIC values separated by a slash (X/X), being the highest value the one that was taken into consideration. Breakpoints of the antibiotics that are contained in EUCAST for *P. aeruginosa* are included in brackets. MIC values ≥ breakpoints are represented as a grey cell. *: PB MICs were elicited in MH II agar. CAZ: ceftazidime, AMK: amikacin, LEV: levofloxacin, CIP: ciprofloxacin, TET: tetracycline, PB: polymyxin B, IPM: imipenem.

**Table S2 | MIC values (µg/ml) to antibiotics of different structural families in *P. aeruginosa* PA14 populations evolved under sub-MIC selective windows of 7 different antibiotics.**

| ANTIBIOTIC | CONCENTRATION (relative to MIC) | REPLICATE (9 days) | CAZ (8) | AMK (16) | LEV (1) | CIP (0,5) | TET | PB* | IPM (4) | ATM (16) | FOF | CHL | ERY |
|---|---|---|---|---|---|---|---|---|---|---|---|---|---|
| **-** | **-** | **PA14** | 0,75 | 2 | 0,125 | 0,047 | 6 | 1 | 1 | 1,5 | 64 | 32 | 48 |
| **-** | **CONTROLS** | **LB1** | 1 | 3 | 0,19 | 0,064 | 6 | 1 | 0,75 | 2 | 48 | 48 | 64 |
| | | **LB2** | 0,75 | 3 | 0,19 | 0,064 | 6 | 1 | 0,75 | 2 | 64 | 48 | 64 |
| | | **LB3** | 0,75 | 3 | 0,19 | 0,064 | 6 | 1 | 0,75 | 2 | 48 | 48 | 64 |
| | | **LB4** | 1 | 3 | 0,19 | 0,064 | 6 | 1,5 | 1 | 2 | 64 | 48 | 64 |
| | | **MHII 1** | 1 | 3 | 0,19 | 0,064 | 6 | 1 | 0,75 | 2 | 48 | 48 | 48 |
| | | **MHII 2** | 0,75 | 3 | 0,19 | 0,064 | 6 | 1 | 1 | 2 | 64 | 32 | 48 |
| | | **MHII 3** | 1 | 3 | 0,19 | 0,064 | 6 | 1,5 | 0,75 | 2 | 48 | 48 | 48 |
| | | **MHII 4** | 1 | 3 | 0,125 | 0,064 | 6 | 1,5 | 1 | 2 | 64 | 48 | 48 |
| **CAZ** | **1/10** | **C9** | - | 12 | 0,38 | 0,094 | 24 | 1 | 0,75 | 4 | 32 | ≥256 | ≥256 |
| | | **C10** | - | 12 | 0,25 | 0,094 | 48 | 1 | 1 | 6 | 64 | ≥256 | ≥256 |
| | | **C11** | - | 8 | 0,38 | 0,094 | 24 | 1 | 1,5 | 6 | 12/24 | ≥256 | ≥256 |
| | | **C12** | - | 12 | 0,38 | 0,064 | 32 | 1,5 | 1 | 6 | 64 | ≥256 | ≥256 |
| | **1/5** | **C13** | - | 16 | 0,75 | 0,19 | 96 | 1 | 1 | 12 | 96 | ≥256 | ≥256 |
| | | **C14** | - | 8 | 0,25 | 0,094 | 24 | 1 | 1/3 | 8 | 48 | ≥256 | ≥256 |
| | | **C15** | - | 6 | 0,5 | 0,064 | 12 | 1 | 8 | 16 | 64 | ≥256 | 192 |
| | | **C16** | - | 6 | 0,38 | 0,094 | 24 | 1 | 0,75 | 12 | 32 | ≥256 | ≥256 |
| | **1/2** | **C17** | - | 1,5/6 | 0,19 | 0,064 | 6 | 0,75 | 1 | 16 | 8/24 | ≥256 | ≥256 |
| | | **C18** | - | 3 | 0,125 | 0,064 | 12 | 0,75 | 16 | 24 | 16 | ≥256 | ≥256 |
| | | **C19** | - | 1,5/12 | 0,19 | 0,047 | 6 | 0,5 | 1,5 | 12 | 8 | ≥256 | ≥256 |

| | | | | | | | | | | | | | |
|---|---|---|---|---|---|---|---|---|---|---|---|---|---|
| | | **C20** | - | 8 | 0,25 | 0,064 | 16 | 0,75 | 1 | 12 | 24 | ≥256 | ≥256 |
| **AMK** | **1/25** | **A9** | 0,75 | - | 0,19 | 0,094 | 6 | 1 | 1 | 2 | 96 | ≥256 | ≥256 |
| | | **A10** | 2 | - | 0,19 | 0,064 | 16 | 1 | 0,75 | 3 | 64 | 128 | ≥256 |
| | | **A12** | 0,75 | - | 0,38 | 0,064 | 8 | 1 | 0,75 | 2 | 64 | 128 | ≥256 |
| | **1/10** | **A13** | 2 | - | 0,38 | 0,094 | 24 | 1 | 1 | 4 | 64 | ≥256 | ≥256 |
| | | **A14** | 2 | - | 0,25 | 0,064 | 24 | 1 | 0,75 | 3 | 256 | ≥256 | ≥256 |
| | | **A15** | 4 | - | 0,5 | 0,094 | 32 | 1,5 | 2 | 8 | 64 | ≥256 | ≥256 |
| | | **A16** | 2 | - | 0,25 | 0,064 | 32 | 1 | 2 | 4 | 32 | ≥256 | ≥256 |
| | **1/5** | **A17** | 3 | - | 0,5 | 0,094 | 32 | 1 | 1 | 3 | 64 | ≥256 | ≥256 |
| | | **A18** | 3 | - | 0,5 | 0,125 | 24 | 1 | 1,5 | 2/8 | 192 | ≥256 | ≥256 |
| | | **A19** | 0,5/2 | - | 0,125/0,5 | 0,47/0,094 | 16 | 1 | 1 | 1,5/4 | 48 | 48 | 192 |
| | | **A20** | 24 | - | 0,5 | 0,19 | 32 | 1 | 2 | 3 | 12 | ≥256 | ≥256 |
| | **1/2** | **A21** | 3 | - | 0,75 | 0,19 | 24 | 1 | 1,5 | 6 | 8 | 128 | ≥256 |
| | | **A22** | 3 | - | 0,5 | 0,125 | 32 | 1 | 2 | 4 | 16 | ≥256 | ≥256 |
| | | **A23** | 2 | - | 0,75 | 0,19 | 48 | 1 | 1,5 | 3 | 16 | ≥256 | ≥256 |
| | | **A24** | 3 | - | 0,5 | 0,19 | 32 | 1,5 | 3 | 4 | 16 | ≥256 | ≥256 |
| | **1/50** | **L5** | 0,75/3 | 4 | - | 0,094 | 12 | 1 | 1 | 1/6 | 64 | 64 | ≥256 |
| | | **L6** | 1,5 | 3/6 | - | 0,064 | 12 | 1 | 0,75 | 2 | 64 | 96 | ≥256 |
| | | **L7** | 1,5 | 8 | - | 0,064 | 24 | 0,75 | 0,75 | 2 | 32 | 192 | ≥256 |
| | **1/25** | **L9** | 0,75/1,5 | 3 | - | 0,19/0,75 | 12 | 1 | 0,75 | 6 | 32 | 96 | ≥256 |
| | | **L10** | 0,5/1,5 | 3/12 | - | 0,75 | 24 | 1 | 0,5 | 2 | 48 | ≥256 | ≥256 |
| | | **L11** | 1 | 2 | - | 0,5 | 16 | 1 | 0,5 | 1,5 | 48 | ≥256 | ≥256 |
| | | **L12** | 1 | 6 | - | 0,75 | 12 | 1 | 0,5 | 1,5 | 64 | ≥256 | ≥256 |
| | **1/10** | **L13** | 1 | 2 | - | 0,75 | 32 | 0,75 | 0,75 | 1 | 48 | ≥256 | 96 |

| | | | | | | | | | | | | | |
|---|---|---|---|---|---|---|---|---|---|---|---|---|---|
| **LEV** | | **L14** | 1 | 2 | - | 0,75 | 16 | 1 | 1,5 | 1,5 | 64 | ≥256 | 128 |
| | | **L15** | 1 | 1,5 | - | 0,75 | 16 | 1 | 1,5 | 1,5 | 48 | ≥256 | 192 |
| | | **L16** | 1 | 1,5 | - | 1 | 24 | 1 | 1 | 1 | 48 | ≥256 | 96 |
| | **1/5** | **L17** | 1 | 2 | - | 1 | 24 | 1 | 0,38 | 2 | 48 | ≥256 | 96 |
| | | **L18** | 1 | 0,5/1,5 | - | 0,75 | 16 | 1 | 0,5 | 1,5 | 32 | ≥256 | 64 |
| | | **L19** | 1 | 2 | - | 1 | 24 | 1 | 0,75 | 1,5 | 48 | ≥256 | 192 |
| | | **L20** | 1 | 2 | - | 1 | 24 | 1 | 0,38 | 1 | 32 | ≥256 | 64 |
| | **1/2** | **L21** | 1,5 | 1,5 | - | 1,5 | 24 | 1 | 1 | 1 | 48 | ≥256 | 64 |
| | | **L22** | 1 | 1,5 | - | 1,5 | 16 | 0,75 | 0,75 | 1,5/2 | 48 | ≥256 | 64 |
| | | **L23** | 1 | 1,5 | - | 2 | 16 | 1 | 0,38 | 1 | 64 | ≥256 | 48 |
| | | **L24** | 1 | 1,5 | - | 12 | 128 | 0,75 | 0,38 | 1 | 48 | ≥256 | 48 |
| **CIP** | **1/100** | **X5** | 2 | 8 | 0,38 | - | 16 | 1 | 1 | 12 | 48 | ≥256 | ≥256 |
| | | **X6** | 1 | 12 | 0,25 | - | 12 | 1 | 1 | 4 | 24 | 192 | ≥256 |
| | | **X7** | 0,75/3 | 3/8 | 0,25 | - | 12 | 1 | 1 | 4 | 24 | ≥256 | ≥256 |
| | | **X8** | 1,5 | 8 | 0,38 | - | 12 | 1 | 1,5 | 3 | 32 | 128 | ≥256 |
| | **1/50** | **X9** | 1,5 | 6 | 2 | - | 24 | 1 | 1 | 2 | 32 | ≥256 | 64 |
| | | **X10** | 1,5 | 4 | 2 | - | 8 | 1 | 1 | 2 | 24 | ≥256 | ≥256 |
| | | **X11** | 1 | 6 | 2 | - | 16 | 1 | 1 | 1/6 | 32 | ≥256 | ≥256 |
| | | **X12** | 1 | 3 | 0,5/2 | - | 8 | 1 | 1 | 1,5/4 | 32 | ≥256 | ≥256 |
| | **1/25** | **X13** | 1,5 | 4 | 2 | - | 16 | 1 | 1 | 6 | 24 | ≥256 | ≥256 |
| | | **X14** | 1 | 2 | 3 | - | 16 | 1 | 0,75 | 0,75 | 24 | ≥256 | 128 |
| | | **X15** | 1 | 2 | 2 | - | 12 | 0,75 | 1 | 2 | 24 | ≥256 | ≥256 |
| | | **X16** | 1 | 3 | 3 | - | 12 | 1,5 | 1,5 | 2 | 64 | ≥256 | ≥256 |
| | **1/10** | **X17** | 1 | 4 | 3 | - | 12 | 1 | 1 | 1 | 16 | ≥256 | ≥256 |

| | | | | | | | | | | | | | |
|---|---|---|---|---|---|---|---|---|---|---|---|---|---|
| | | **X18** | 1 | 6 | 4 | - | 16 | 1 | 0,75 | 2 | 24 | ≥256 | ≥256 |
| | | **X19** | 2 | 4 | 4 | - | 16 | 1 | 0,75 | 1,5/4 | 24 | ≥256 | 128 |
| | | **X20** | 1,5 | 4 | 4 | - | 24 | 1 | 0,75 | 2 | 24 | ≥256 | 128 |
| | **1/5** | **X21** | 0,75 | 2 | 24 | - | 48 | 1,5 | 2 | 1 | 16 | ≥256 | 192 |
| | | **X22** | 1,5 | 2 | 6 | - | 24 | 1,5 | 1,5 | 1 | 24 | ≥256 | 96 |
| | | **X23** | 1 | 1,5 | 12 | - | 64 | 1 | 2 | 1 | 24 | ≥256 | 128 |
| | | **X24** | 0,75 | 1,5 | 12 | - | 32 | 1 | 0,75 | 1 | 16 | ≥256 | 128 |
| | **1/2** | **X25** | 1 | 3 | ≥32 | - | 96 | 1 | 0,75 | 0,75 | 8 | ≥256 | ≥256 |
| | | **X26** | 0,75 | 1,5 | ≥32 | - | 12 | 1 | 2 | 1,5 | 24 | ≥256 | 64 |
| | | **X27** | 0,75 | 1,5 | ≥32 | - | 64 | 1 | 0,75 | 0,75 | 16 | ≥256 | ≥256 |
| | | **X28** | 0,75 | 1 | ≥32 | - | 16 | 1 | 1,5 | 1 | 16 | ≥256 | 64 |
| **TET** | **1/10** | **T13** | 3 | 8 | 0,5 | 0,125 | - | 1 | 1 | 6 | 64 | ≥256 | ≥256 |
| | | **T14** | 3 | 12 | 0,38 | 0,125 | - | 1 | 0,75 | 6 | 96 | ≥256 | ≥256 |
| | | **T15** | 4 | 12 | 0,5 | 0,094 | - | 1 | 0,75 | 4 | 64 | ≥256 | ≥256 |
| | | **T16** | 3 | 12 | 0,25 | 0,064 | - | 1,5 | 1 | 4 | 96 | ≥256 | ≥256 |
| | **1/5** | **T17** | 2 | 12 | 0,38 | 0,094 | - | 1 | 1 | 6 | 96 | 192 | ≥256 |
| | | **T18** | 3 | 8 | 0,38 | 0,094 | - | 1 | 1 | 6 | 16 | 192 | ≥256 |
| | | **T19** | 3 | 12 | 0,38 | 0,094/0,5 | - | 1,5 | 0,75 | 6 | 64 | ≥256 | ≥256 |
| | | **T20** | 3 | 8 | 0,38 | 0,125 | - | 1 | 1 | 4 | 12/48 | ≥256 | ≥256 |
| | **1/2** | **T21** | 4 | 6 | 0,25/2 | 1 | - | 1 | 1 | 4 | 12/48 | 96 | ≥256 |
| | | **T22** | 8 | 6 | 1 | 0,125 | - | 1 | 1,5 | 24 | 16/64 | ≥256 | ≥256 |
| | | **T23** | 4 | 8 | 0,75 | 0,094/0,5 | - | 1 | 1,5 | 4/16 | 12/32 | ≥256 | ≥256 |
| | | **T24** | 3 | 12 | 0,25/1,5 | 0,094 | - | 1,5 | 1,5 | 6 | 16/48 | 128 | ≥256 |
| | **1/2** | **P10** | 1,5 | 4 | 0,38 | 0,064 | 12 | - | 1,5/2 | 1,5 | 48 | 32 | ≥256 |

| | | | | | | | | | | | | | |
|---|---|---|---|---|---|---|---|---|---|---|---|---|---|
| **PB** | | **P11** | 0,75 | 6/24 | 0,38 | 0,125 | 12 | - | 1 | 1 | 32 | 24 | ≥256 |
| **IPM** | **1/2** | **I9** | 0,75/3 | 3/8 | 0,38 | 0,064 | 12 | 1 | - | 6 | 64 | 128 | ≥256 |
| | | **I10** | 1,5 | 3/8 | 0,38 | 0,064 | 12 | 0,5 | - | 2 | 64 | 64 | ≥256 |
| | | **I11** | 0,75/2 | 2/8 | 0,38 | 0,094 | 12 | 1 | - | 4 | 64 | 64 | ≥256 |

All MICs are obtained by E-test strips. Double inhibition halos are shown as two MIC values separated by a slash (X/X), being the highest value the one that is taken into consideration. Breakpoints of the antibiotics that are included in EUCAST for *P. aeruginosa* are included in brackets. MIC values ≥ breakpoints are represented as a grey cell. LB1-4 and MH II 1-4 refer to control populations evolved in absence of drug (in LB and MH II medium, respectively), being their MICs at 3 and 6 days identical to the ones at 9 days. *: PB MICs were elicited in MH II agar. CAZ: ceftazidime, AMK: amikacin, LEV: levofloxacin, CIP: ciprofloxacin, TET: tetracycline, PB: polymyxin B, IPM: imipenem, ATM: aztreonam, FOF: Fosfomycin, CHL: chloramphenicol, ERY: erythromycin.

**Table S3 | MIC values (µg/ml) to antibiotics of different structural families in presence and absence of EPI PAβN in *P. aeruginosa* PA14 populations evolved under sub-MIC selective windows of 7 different antibiotics.**

| ANTIBIOTIC | CONCENTRATION (relative to MIC) | REPLICATE | CAZ | | AMK | | LEV | | CIP | | TET | | PB | | IPM | | CHL | | ERY | |
|---|---|---|---|---|---|---|---|---|---|---|---|---|---|---|---|---|---|---|---|---|
| | | | MH | PAβN | MH | PAβN | MH | PAβN | MH | PAβN | MH | PAβN | MH | PAβN | MH | PAβN | MH | PAβN | MH | PAβN |
| - | - | **PA14** | 0,75 | 0,75 | 2 | 2 | 0,125 | 0,023 | 0,047 | 0,032 | 6 | 6 | 1 | 1 | 1 | 1 | 32 | 4 | 48 | 24 |
| **CAZ** | **1/10** | **C9** | 2 | 1 | 12 | 12 | 0,38 | 0,016 | - | - | 24 | 6 | - | - | - | - | - | - | - | - |
| | | **C10** | 2 | 1 | 12 | 12 | 0,25 | 0,023 | - | - | 48 | 6 | - | - | - | - | - | - | - | - |
| | | **C11** | 2 | 0,25 | 8 | 8 | 0,38 | 0,016 | - | - | 24 | 6 | - | - | - | - | - | - | - | - |
| | | **C12** | 3 | 0,25 | 12 | 16 | 0,38 | 0,016 | - | - | 32 | 8 | - | - | - | - | - | - | - | - |
| | **1/5** | **C13** | 4 | 1 | 16 | 16 | 0,75 | 0,032 | - | - | 96 | 16 | - | - | - | - | - | - | - | - |
| | | **C14** | 2 | 0,38 | 8 | 8 | 0,25 | 0,023 | - | - | 24 | 6 | - | - | - | - | - | - | - | - |
| | | **C15** | 8 | 3 | 6 | 1/6 | 0,5 | 0,016 | - | - | 12 | 4 | - | - | - | - | - | - | - | - |
| | | **C16** | 4 | 0,75 | 6 | 6 | 0,38 | 0,023 | - | - | 24 | 6 | - | - | - | - | - | - | - | - |
| | **1/2** | **C17** | 6 | X | 6 | X | 0,9 | X | - | - | 6 | X | - | - | - | - | - | - | - | - |
| | | **C18** | 12 | X | 3 | X | 0,125 | X | - | - | 12 | X | - | - | - | - | - | - | - | - |
| | | **C19** | 8 | X | 1/12 | X | 0,19 | X | - | - | 6 | X | - | - | - | - | - | - | - | - |
| | | **C20** | 6 | X | 8 | X | 0,25 | X | - | - | 16 | X | - | - | - | - | - | - | - | - |
| | **1/25** | **A9** | 0,75 | 0,75 | 4 | 3 | 0,19 | 0,047 | - | - | 6 | 6 | - | - | - | - | ≥256 | 4 | - | - |
| | | **A10** | 2 | 1 | 2/4 | 3 | 0,19 | 0,023 | - | - | 16 | 6 | - | - | - | - | 128 | 4 | - | - |
| | | **A12** | 0,75 | 0,75 | 6 | 6 | 0,38 | 0,032 | - | - | 8 | 6 | - | - | - | - | 128 | 6 | - | - |
| | **1/10** | **A13** | 2 | 1 | 8 | 8 | 0,38 | 0,032 | - | - | 24 | 12 | - | - | - | - | ≥256 | 4 | - | - |
| | | **A14** | 2 | 1 | 16 | 3/12 | 0,25 | 0,032 | - | - | 24 | 8 | - | - | - | - | ≥256 | 6 | - | - |
| | | **A15** | 4 | 1 | 12 | 12 | 0,5 | 0,016 | - | - | 32 | 6 | - | - | - | - | ≥256 | 3 | - | - |

| | | | | | | | | | | | | | | | | | | | | |
|---|---|---|---|---|---|---|---|---|---|---|---|---|---|---|---|---|---|---|---|---|
| **AMK** | | **A16** | 2 | 1 | 12 | 12 | 0,25 | 0,032 | - | - | 32 | 8 | - | - | - | - | ≥256 | 6 | - | - |
| | **1/5** | **A17** | 3 | 1 | 24 | 24 | 0,5 | 0,047 | - | - | 32 | 8 | - | - | - | - | ≥256 | 4 | - | - |
| | | **A18** | 3 | 1 | 16 | 8/24 | 0,5 | 0,032 | - | - | 24 | 6 | - | - | - | - | ≥256 | 3 | - | - |
| | | **A19** | 0,5/2 | 0,75 | 8 | 24 | 0,5 | 0,023 | - | - | 16 | 6 | - | - | - | - | 48 | 1 | - | - |
| | | **A20** | 24 | 0,75 | 24 | 32 | 0,5 | 0,023 | - | - | 32 | 6 | - | - | - | - | ≥256 | 2 | - | - |
| | **1/2** | **A21** | 3 | 4 | 24 | 24 | 0,75 | 0,047 | - | - | 24 | 24 | - | - | - | - | 128 | ≥256 | - | - |
| | | **A22** | 3 | 0,5 | 24 | 32 | 0,5 | 0,023 | - | - | 32 | 6 | - | - | - | - | ≥256 | 2 | - | - |
| | | **A23** | 2 | 0,75 | 24 | 48 | 0,75 | 0,023 | - | - | 48 | 8 | - | - | - | - | ≥256 | 2 | - | - |
| | | **A24** | 3 | 1 | 24 | 32 | 0,5 | 0,023 | - | - | 32 | 6 | - | - | - | - | ≥256 | 2 | - | - |
| **LEV** | **1/50** | **L5** | 0,75/3 | 0,75 | 4 | 3 | 0,25/0,5 | 0,032 | - | - | 12 | 4 | - | - | - | - | 64 | 6 | ≥256 | 3/24 |
| | | **L6** | 1,5 | 0,75 | 3/6 | 4 | 0,19/0,5 | 0,032 | - | - | 12 | 6 | - | - | - | - | 96 | 4 | ≥256 | 16 |
| | | **L7** | 1,5 | 0,75 | 8 | 1/6 | 0,19/2 | 0,023 | - | - | 24 | 6 | - | - | - | - | 192 | 4 | ≥256 | 4/24 |
| | **1/25** | **L9** | - | - | - | - | 2 | 0,094 | - | - | 12 | 12 | - | - | - | - | 96 | 64 | ≥256 | ≥256 |
| | | **L10** | - | - | - | - | 0,75/3 | 0,094 | - | - | 24 | 16 | - | - | - | - | ≥256 | 32 | ≥256 | 96 |
| | | **L11** | - | - | - | - | 3 | 0,094 | - | - | 16 | 16 | - | - | - | - | ≥256 | ≥256 | ≥256 | ≥256 |
| | | **L12** | - | - | - | - | 4 | 0,094 | - | - | 12 | 12 | - | - | - | - | ≥256 | 48 | ≥256 | ≥256 |
| | **1/10** | **L13** | - | - | - | - | 3 | 0,125 | - | - | 32 | 12 | - | - | - | - | ≥256 | 128 | 96 | 48 |
| | | **L14** | - | - | - | - | 3 | 0,125 | - | - | 16 | 12 | - | - | - | - | ≥256 | 128 | 128 | 64 |
| | | **L15** | - | - | - | - | 3 | 0,064 | - | - | 16 | 12 | - | - | - | - | ≥256 | 96 | 192 | 48 |
| | | **L16** | - | - | - | - | 4 | 0,19 | - | - | 24 | 16 | - | - | - | - | ≥256 | ≥256 | 96 | 32 |
| | **1/5** | **L17** | - | - | - | - | 3 | 0,125 | - | - | 24 | 16 | - | - | - | - | ≥256 | 128 | 96 | 48 |
| | | **L18** | - | - | - | - | 4 | 0,125 | - | - | 16 | 12 | - | - | - | - | ≥256 | 96 | 64 | 48 |
| | | **L19** | - | - | - | - | 3 | 0,125 | - | - | 24 | 16 | - | - | - | - | ≥256 | ≥256 | 192 | 64 |
| | | **L20** | - | - | - | - | 4 | 0,125 | - | - | 24 | 8 | - | - | - | - | ≥256 | 64 | 64 | 24 |

| | | | | | | | | | | | | | | | | | | | | |
|---|---|---|---|---|---|---|---|---|---|---|---|---|---|---|---|---|---|---|---|---|
| | **1/2** | **L21** | - | - | - | - | 8 | 0,125 | - | - | 24 | 3 | - | - | - | - | ≥256 | 96 | 64 | 16 |
| | | **L22** | - | - | - | - | 6 | 0,125 | - | - | 16 | 4 | - | - | - | - | ≥256 | 128 | 64 | 24 |
| | | **L23** | - | - | - | - | 12 | 0,19 | - | - | 16 | 3 | - | - | - | - | ≥256 | 96 | 48 | 12 |
| | | **L24** | - | - | - | - | ≥32 | 0,5 | - | - | 128 | 3 | - | - | - | - | ≥256 | 96 | 48 | 6 |
| **CIP** | **1/100** | **X5** | - | - | 8 | 6 | - | - | 0,125 | 0,032 | 16 | 16 | - | - | - | - | ≥256 | 4 | - | - |
| | | **X6** | - | - | 12 | 4 | - | - | 0,125 | 0,047 | 12 | 12 | - | - | - | - | 192 | 6 | - | - |
| | | **X7** | - | - | 3/8 | 6 | - | - | 0,094 | 0,032 | 12 | 12 | - | - | - | - | ≥256 | 64 | - | - |
| | | **X8** | - | - | 8 | 3 | - | - | 0,094 | 0,032 | 12 | 12 | - | - | - | - | 128 | 6 | - | - |
| | **1/50** | **X9** | - | - | 6 | 6 | - | - | 0,75 | 0,19 | 24 | 24 | - | - | - | - | ≥256 | 24 | - | - |
| | | **X10** | - | - | 4 | 3 | - | - | 0,75 | 0,19 | 8 | 6 | - | - | - | - | ≥256 | 48 | - | - |
| | | **X11** | - | - | 6 | 6 | - | - | 0,75 | 0,19 | 16 | 12 | - | - | - | - | ≥256 | 64 | - | - |
| | | **X12** | - | - | 3 | 3 | - | - | 0,75 | 0,094 | 8 | 8 | - | - | - | - | ≥256 | 32 | - | - |
| | **1/25** | **X13** | - | - | 4 | 3 | - | - | 1 | 0,19 | 16 | 12 | - | - | - | - | ≥256 | 48 | - | - |
| | | **X14** | - | - | 2 | 1,5 | - | - | 1 | 0,19 | 16 | 16 | - | - | - | - | ≥256 | 192 | - | - |
| | | **X15** | - | - | 2 | 2 | - | - | 0,75 | 0,19 | 12 | 12 | - | - | - | - | ≥256 | 32 | - | - |
| | | **X16** | - | - | 3 | 3 | - | - | 1,5 | 0,19 | 12 | 16 | - | - | - | - | ≥256 | 64 | - | - |
| | **1/10** | **X17** | - | - | 4 | 3 | - | - | 1,5 | 0,25 | 12 | 12 | - | - | - | - | ≥256 | 96 | - | - |
| | | **X18** | - | - | 6 | 6 | - | - | 1,5 | 0,25 | 16 | 16 | - | - | - | - | ≥256 | ≥256 | - | - |
| | | **X19** | - | - | 4 | 2 | - | - | 1 | 0,19 | 16 | 12 | - | - | - | - | ≥256 | ≥256 | - | - |
| | | **X20** | - | - | 4 | 2 | - | - | 1,5 | 0,25 | 24 | 24 | - | - | - | - | ≥256 | 96 | - | - |
| | **1/5** | **X21** | - | - | 2 | 1,5 | - | - | 16 | 0,75 | 48 | 24 | - | - | - | - | ≥256 | 64 | - | - |
| | | **X22** | - | - | 2 | 2 | - | - | 3 | 0,125 | 24 | 16 | - | - | - | - | ≥256 | 128 | - | - |
| | | **X23** | - | - | 1,5 | 1,5 | - | - | 3 | 0,25 | 64 | 16 | - | - | - | - | ≥256 | 128 | - | - |
| | | **X24** | - | - | 1,5 | 1 | - | - | 4 | 0,19 | 32 | 24 | - | - | - | - | ≥256 | ≥256 | - | - |
| | | **X25** | - | - | 3 | 0,75 | - | - | ≥32 | 0,38 | 96 | 6 | - | - | - | - | ≥256 | 32 | - | - |

| | | | | | | | | | | | | | | | | | | | | |
|---|---|---|---|---|---|---|---|---|---|---|---|---|---|---|---|---|---|---|---|---|
| | **1/2** | **X26** | - | - | 1,5 | 1,5 | - | - | 6 | 0,25 | 12 | 12 | - | - | - | - | ≥256 | 64 | - | - |
| | | **X27** | - | - | 1,5 | 1 | - | - | 12 | 0,25 | 64 | 6 | - | - | - | - | ≥256 | 64 | - | - |
| | | **X28** | - | - | 1 | 1,5 | - | - | 6 | 0,5 | 16 | 12 | - | - | - | - | ≥256 | 64 | - | - |
| **TET** | **1/10** | **T13** | 3 | 1,5 | 8 | 1,5/8 | 0,5 | 0,032 | - | - | 12 | 4 | - | - | - | - | - | - | - | - |
| | | **T14** | 3 | 1 | 12 | 12 | 0,38 | 0,023 | - | - | 16 | 6 | - | - | - | - | - | - | - | - |
| | | **T15** | 4 | 1 | 1/12 | 1/12 | 0,5 | 0,023 | - | - | 24 | 4 | - | - | - | - | - | - | - | - |
| | | **T16** | 3 | 1 | 12 | 3/12 | 0,25 | 0,032 | - | - | 16 | 6 | - | - | - | - | - | - | - | - |
| | **1/5** | **T17** | 2 | 0,75 | 12 | 4/24 | 0,38 | 0,032 | - | - | 24 | 6 | - | - | - | - | - | - | - | - |
| | | **T18** | 3 | 0,5 | 8 | 8 | 0,38 | 0,023 | - | - | 24 | 6 | - | - | - | - | - | - | - | - |
| | | **T19** | 3 | 1 | 12 | 12 | 0,38 | 0,023 | - | - | 16 | 4 | - | - | - | - | - | - | - | - |
| | | **T20** | 3 | 0,75 | 8 | 6 | 0,38 | 0,016 | - | - | 24 | 6 | - | - | - | - | - | - | - | - |
| | **1/2** | **T21** | 4 | 0,75 | 6 | 6 | 0,25/2 | 0,094 | - | - | 24 | 4 | - | - | - | - | - | - | - | - |
| | | **T22** | 8 | 3 | 6 | 4 | 1 | 0,023 | - | - | 48 | 8 | - | - | - | - | - | - | - | - |
| | | **T23** | 4 | 1 | 8 | 8 | 0,75 | 0,064 | - | - | 24 | 4 | - | - | - | - | - | - | - | - |
| | | **T24** | 3 | 0,5 | 12 | 12 | 1,5 | 0,047 | - | - | 24 | 6 | - | - | - | - | - | - | - | - |
| **PB** | **1/2** | **P10** | - | - | 4 | 3 | 0,38 | 0,25 | - | - | 12 | 12 | 2 | 2 | - | - | - | - | - | - |
| | | **P11** | - | - | 6/24 | 6 | 0,38 | 0,38 | - | - | 12 | 12 | 6 | 6 | - | - | - | - | - | - |
| **IPM** | **1/2** | **I9** | 0,75/3 | 2 | 3/8 | 3/8 | 0,38 | 0,032 | - | - | 12 | 12 | - | - | 2 | 0,5 | - | - | - | - |
| | | **I10** | 1,5 | 1 | 3/8 | 8 | 0,38 | 0,023 | - | - | 12 | 12 | - | - | 2 | 0,5 | - | - | - | - |
| | | **I11** | 0,75/2 | 1,5 | 2/8 | 2/8 | 0,38 | 0,023 | - | - | 12 | 12 | - | - | 2 | 0,75 | - | - | - | - |

All MICs are obtained by E-test strips, in absence and in presence of 25 µg/ml of EPI PAβN. Double inhibition halos are shown as two MIC values separated by a slash (X/X), being the highest value the one that was taken into consideration. “X” indicates the replicates that were unable to grow in presence of PAβN, and “-“ the MICs that were not determined in presence of this EPI. CAZ: ceftazidime, AMK:

amikacin, LEV: levofloxacin, CIP: ciprofloxacin, TET: tetracycline, PB: polymyxin B, IPM: imipenem, CHL: chloramphenicol, ERY: erythromycin.
